# Meta-Analysis of cortical inhibitory interneurons markers landscape and their performances in scRNA-seq studies

**DOI:** 10.1101/2021.11.03.467049

**Authors:** Lorenzo Martini, Roberta Bardini, Stefano Di Carlo

## Abstract

The mammalian cortex contains a great variety of neuronal cells. In particular, GABAergic interneurons, which play a major role in neuronal circuit function, exhibit an extraordinary diversity of cell types. In this regard, single-cell RNA-seq analysis is crucial to study cellular heterogeneity. To identify and analyze rare cell types, it is necessary to reliably label cells through known markers. In this way, all the related studies are dependent on the quality of the employed marker genes. Therefore, in this work, we investigate how a set of chosen inhibitory interneurons markers perform. The gene set consists of both immunohistochemistry-derived genes and single-cell RNA-seq taxonomy ones. We employed various human and mouse datasets of the brain cortex, consequently processed with the Monocle3 pipeline. We defined metrics based on the relations between unsupervised cluster results and the marker expression. Specifically, we calculated the specificity, the fraction of cells expressing, and some metrics derived from decision tree analysis like entropy gain and impurity reduction. The results highlighted the strong reliability of some markers but also the low quality of others. More interestingly, though, a correlation emerges between the general performances of the genes set and the experimental quality of the datasets. Therefore, the proposed method allows evaluating the quality of a dataset in relation to its reliability regarding the inhibitory interneurons cellular heterogeneity study.

## I. Introduction

The cerebral cortex is one of the most complex systems in biology, with an extraordinary variety of specialized cells forming the neuronal circuit at the base of the brain functions [1]. It is composed primarily of neuronal and non-neuronal cells. Neuronal cells are mainly divided into excitatory (or glutamatergic) and inhibitory (or GABAergic) neurons. Excitatory neurons comprise the majority of cells and connect distal areas of the cortex. Inhibitory neurons, primarily composed of interneurons, connect locally to excitatory neurons and inhibit the transmission in neuronal circuits. In particular, GABAergic interneurons show several subtypes, which differ in morphology, functionality, neurochemical and physiological properties. The deep and complete understanding and taxonomy of inhibitory interneurons is still an open problem in biology [2].

To help with this challenge, the employment of singlecell RNA sequencing (scRNA-seq) data is important. ScRNA-seq allows the investigation of the gene-expression profile with single-cell resolution, enabling the study of cellular heterogeneity at the transcriptional level [4][5]. The scRNA-seq analysis typically involves unsupervised clustering algorithms, which cluster cells of a dataset into groups based on mathematical and topology-based algorithms. However, these clusters, and more importantly, the cells they contain, have no biological identities. To label them, one usually performs differential expression analysis or investigates the expression of cell-type-related marker genes. In both cases, the discriminative ability and reliability of the selected marker genes are crucial for heterogeneity studies, especially regarding the rare cell types identification.

This paper focuses on GABAergic interneurons subtypes identification. It aims at investigating the landscape of biomarkers commonly employed for their analysis. One of the main contributions is the analysis of the marker reliability when applied to the scRNA-seq analysis. The analysis takes into consideration literature-based markers from immunohistochemistry studies and single-cell taxonomy analysis. Moreover, the paper defines metrics to highlight and evaluate the marker’s performances to enable quantitative comparison. These metrics have been defined to be general and easy to apply, so one could employ them to investigate other biomarkers for various cell type populations.

## II. Materials and Methods

### A. Literature Markers study

We started with an extensive literature review to understand how the state-of-the-art defines the subtypes of the cortical GABAergic interneurons. We initially focused on various studies involving this type of cells, including immunohistochemistry (Figure 1a) and scRNA-seq (Figure 1b) analyses. The result is an increasingly complicated landscape of markers and subgroups. Here we give an overview of their classification, considering different levels of hierarchy. First, all inhibitory neurons are characterized by using *γ*-aminobutyric acid as a neurotransmitter strictly related to the enzyme glutamic acid decarboxylase (GAD), encoded by the genes *Gad1* and *Gad2* [6]. These two genes are notoriously employed to identify inhibitory neurons as a whole and are broadly accepted as their markers. Then cortical interneurons are divided into two categories, dependent on their anatomical origin in the developmental cortex [3][7]. The first one is the *medial ganglionic eminence* (MGE) derived cells from which the majority (about 70% [8]) of the cortical interneurons derives. The remaining cells are *central ganglionic eminence* (CGE)-derived. Regarding, this subdivision, we identified, form immunohistochemistry the gene 5-Hydroxytryptamine Receptor 3A *(Htr3a)* linked to CGE-derived cells, and from scRNA-seq studies the genes Adenosine Deaminase RNA Specific B2 *(Adrab2)* and LIM Homeobox 6 *(Lhx6)* respectively for MGE and CGE derived cells. These two subgroups give then origin to a constellation of subtypes [8][1]. For this research, we investigate them using gene expression. The MGE-derived cells are classified into two non-overlapping subtypes, the *parvalbumin-expressing* (PVALB), and the *somatostatin-expressing* (SST) cells, named after their characterizing genes *Pvalb* and *Sst*. Our investigation of the immunohistochemistry landscape and single-cell taxonomy leads to a complex landscape of genes, whose expression defines subclasses of interneurons overlapping with each other (Figure 1a) [8][3][9]. Therefore, after this preliminary analysis, we decided to divide the remaining CGE-derived cells, into two subgroups. The *vasoactive intestinal peptide-expressing* (VIP) cells related to the *Vip* gene, and *non-Vip-expressing*(LAMP) cells with the Lysosomal Associated Membrane Protein Family Member 5 *(Lamp5)* gene as a marker. Overall, Figure 1 recapitulates the division and the related markers of the cortical inhibitory interneurons. Based on this preliminary analysis, we decided to consider a division into four subclasses, namely PVALB, SST, VIP, and LAMP, based on the expression of the genes *Pvalb, Sst, Vip, Lamp5.* Two classes (i.e., PVALB and SST) are associated with CGE-derived cells, and two classes (i.e., VIP and LAMP) are associated with MGE-derived cells. We considered a collection of 9 marker genes (i.e., *Gad1, Gad2, Adrb2, Htr3a, Lhx6, Pvalb, Sst, Vip,* and *Lamp5)* for which we accurately calculated the defined metrics to understand their capability to discriminate different classes of cells reliably.

**Fig. 1.**
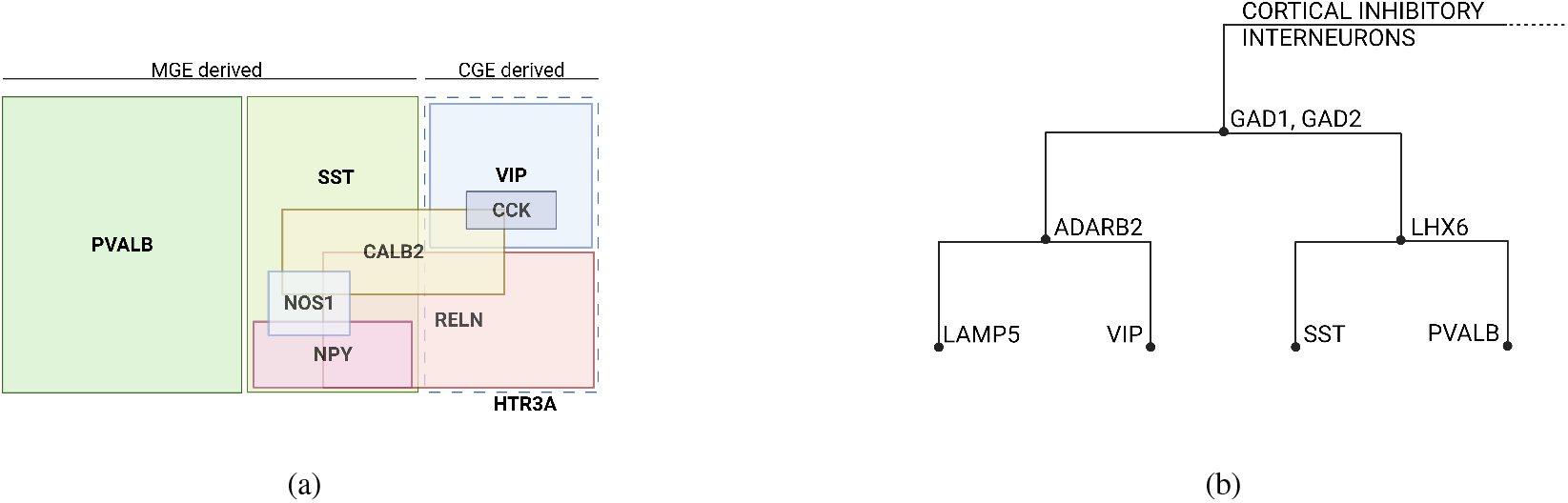
Cortical inhibitory interneurons biomarkers are derived from the analysis of the literature. (a) Biomarkers identified using immunohistochemistry studies (adapted from [3]). (b) Taxonomy and related markers derived from scRNA-seq analysis studies.

### B. Data collection and availability

This work aims to study the overall performance of some marker genes within the context of scRNA-seq analysis. Therefore we need a collection of different datasets to perform a significant meta-analysis able to characterize an average single-cell experiment. Since the previous discussion on cortical inhibitory interneuron diversity is valid for the general mammalian brain cortex, we included datasets from *Mus Musculus* and *Homo Sapiens* samples and examined homologous markers between the two species. To account for possible differences, we refer to the Hodge et al. study [10]. All the samples are from comprehensive cortex studies or specific anatomical and functional cortex regions. We collected and processed a total of 9 datasets: three from *Homo Sapiens* and six from *Mus Musculus*. The two main sources for the acquisition were the Allen Brain Map database [11] of the Allen Institute and the Single Cell Portal [12]. Specifically, four datasets come from Allen (i.e., Allen Human ACC [13], Allen Human MTG [14], Allen Human VISP [13], and Allen Mouse Cortex [15]), four from the Single Cell Portal (i.e., SCP2 mouse AUD [16], SCP3 mouse ORB [17], SCP5 mouse Visp [18], and SCP6 mouse MOP [19]), and a dataset from a multimodal technique experiment (SNARE mouse cortex, GEO accession number: GSE126074 [20]).

### C. Datasets processing

To process the scRNA-seq datasets, we employed the well-known pipeline Monocle3 [21] implemented in R [22]. In this way, we guarantee to mimic the workflow typically employed in single-cell data analysis [5]. For each dataset, we followed the standard workflow illustrated in their vignette []. In particular, one must pay attention to the clustering resolution. Since we are working with four types of inhibitory interneurons (i.e., PVALB, SST, VIP, and LAMP), we decided to work with a final clustering division including four clusters. This enabled us to qualitatively examine the expression of the selected markers and their ability to identify the correct clusters of inhibitory interneurons. This step is important since it defines the cell type classification on which we measured the performance of the related marker genes. Moreover, besides *Gad1* and *Gad2* that are global markers whose metrics can be computed taking into account all the cells in the datasets, for the remaining markers, only cells labeled with a specific group have been considered in the metric calculation.

The full set of R scripts required to reproduce the proposed meta-analysis are deposited in an open repository on GitHub: https://github.com/sysbio-polito/cRNA-seq-interneurons-markers-analysis. Interested readers can refer to the documentation available in the repository for additional technical details.

### D. Metrics definition

To evaluate the performance of the marker genes, we first need to define a set of metrics designed for this task. Good metrics should reflect how a gene is strictly related to a particular group of cells and how good a feature is for the cell classification. Regardless, we need a ground truth classification to calculate the metrics. For this purpose, we considered the unsupervised clusters obtained from the dataset processing. Specifically, we labeled the clusters with the four defined subtypes (i.e., PVALB, SST, VIP, and LAMP), qualitatively assessing the overall expression of the related genes. With this in mind, we defined four metrics.

First of all, we want to measure how much a marker is associated with a cell type. The idea is that a perfect cell type marker is expressed by all the cells it identifies and only by them. For this purpose, we established two correlated metrics: the *fraction expressing* and the *specificity*. The *fraction expressing* is simply the percentage of cells expressing the marker in the cluster. The *specificity* instead indicates how much the marker expression is specific to a cluster. In particular, an optimal marker is expressed only inside the related cluster. We applied a method based on the Jensen-Shannon divergence [23], also used in Monocle3 algorithms to calculate the specificity during differential analysis [24]. Of course, these two metrics are not optimal if employed alone, but it is better to look at their correlation.

The other two metrics, instead, are an evaluation of the informativeness of the markers. Specifically, we treat the cluster classification problem like a decision tree problem, and we try to understand how much the features are informative for the clusters. Therefore, we initially take the cluster classification as a training set, breaking it into subgroups based on some features. In this case, the features are the expression levels of the markers in each cell. Given the resulting partitioning, one can evaluate the classification power of the marker with some well-known metrics: the *Information Gain* [23] and the *Impurity Reduction* [25]. The first is based on Shannon Entropy (H), the second is based on Gini Index (G). The Shannon Entropy (1) measures the amount of uncertainty of a random variable with a probability distribution [23]. Its value ranges between 0 in the case of a certain event (i.e., classification with all equal values) and 1 in the case of a completely random event.

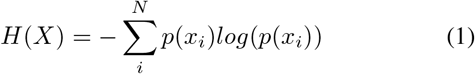

In our case, given a classification X (i.e., the list of all cells’ labels) with N possible classes (i.e., the number of subtypes we are considering), *p*(*x_i_*) represents the probability of the i-th class *x_i_*. Thus, the sum is over all the N classes. The Gini Index (2) calculates the probability of a selected element to be incorrectly classified when randomly labeled [25]. Like H, it assumes values between 0 (i.e., purity of classification) and 1 (i.e., random distribution).

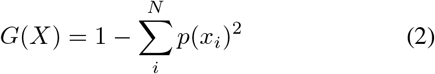

*p*(*x_i_*) and *x_i_* have the same meaning as for the Shannon Entropy. These two metrics are similar in what they estimate, and one could employ only one of the two. However, it is interesting to calculate both since we do not know which one fits better our classification problem. Anyway, they are optimal to evaluate the performance in a classification scenario, a relevant task in labeling a scRNA-seq dataset. It is important to highlight that all the metrics presented here are independent of the particular cellular heterogeneity study but can be applied to examine any set of markers.

### E. Metrics calculation

After the definition of the metrics, we explain how we calculated them for each dataset. Given a processed dataset with the interneurons clusters identified, we calculated the four metrics for the list of genes on three different levels of resolution as summarized in Figure 2.

**Fig. 2.**
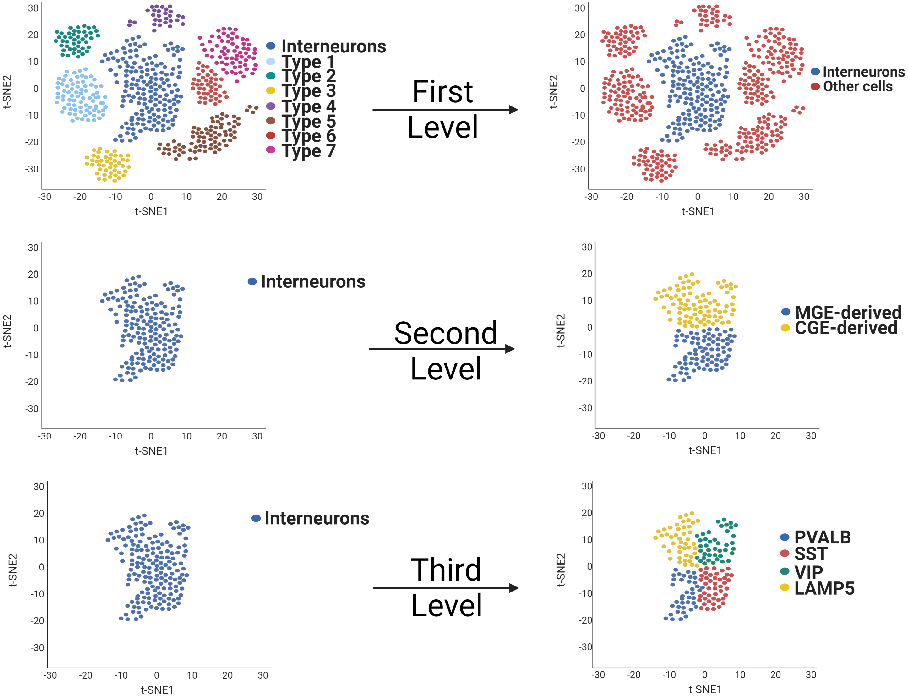
Schematic representation of the cell clustering for metrics calculation. At first, for the markers *Gad1* and *Gad2,* a heterogeneous population of cells from the datasets (heterogeneity is depicted in an abstract way with different colors) was considered. Cells are split into two clusters: interneurons and other cells. Only interneurons are considered and analyzed at the second and third levels, considering respectively two and four subgroups.

We started with the genes *Gad1* and *Gad2,* and we evaluated their performance to discriminate the interneurons from all the other cells (First Level in Figure 2). Therefore, we considered all the identified interneurons as a single cluster. This result represents the starting point for the following levels that aim to split the interneurons’ population into different sets of clusters.

At the next level (Second Level in Figure 2), we evaluate the metrics for the genes able to discriminate between MGE and CGE derived cells starting from the set of interneuron cells identified at the first level and dividing them into two clusters labeled from the expression inspection. All the metrics were calculated based on this partition. Finally, we applied the same reasoning to the next level of classification (Third Level in Figure 2), where we considered the cells grouped into four clusters.

The *fraction expressing* is the fraction of cells in a given cluster expressing the considered marker. It is determined by counting the cells with a marker expression greater than zero. One could introduce a minimum expression threshold to detect the cells, but this would require a meaningful and reliable way to set this level. The *specificity,* as previously mentioned, is based on the Jensen-Shannon divergence. It measures how specific the marker’s expression is to the cell group. The resulting value ranges between 0 and 1. While we measured these two metrics, we also calculated the *mean expression value* and the *marker score*. The marker score is just the product of specificity and fraction expressing. Thus, it is a single value that embeds the two metrics. However, due to its simple nature, it could hide some relevant information if examined alone. The mean expression is the average expression value of the gene in the cluster. It is useful to shed light on the expression levels in a cell type. Moreover, it adds an information layer to the marker evaluation, meaning that there could be a relation between the performance of a marker and its expression. Thus, both the mean expression and the marker score are additional information extending the marker performance evaluation. Information Gain and Impurity Reduction calculations are analogous, besides the actual mathematical formula employed. We started by calculating the Entropy and Gini Index of the cluster classification. Then, given a marker, we divided the cells into two groups: those that do and do not express it. For each subgroup, we calculated the two functions again. IG and IR were therefore calculated as the subtraction between before and after. This means that a good marker, which correctly classifies the cells, has a high Information Gain. The same reasoning is also valid for Impurity Reduction, which indicates how pure the classification is after the split.

## III. Results

We present here the results of the proposed analysis applied to the selected set of markers on all considered datasets.

We start with the analysis of the *specificity* and *fraction expressing*. As previously mentioned, individually investigating the two metrics is not optimal since a single value does not give a complete insight into the performance of a marker. For example, a gene with very high *specificity* but low *fraction expressing* implies that only a small population in a specific cluster expresses it, which is not optimal when looking for a good marker. For this reason, we look at them concurrently. The best and easiest method is to visualize them on a plot of *specificity* vs. *fraction expressing*, as reported in Figure 3. All markers are labeled with their name and colored based on the origin dataset. The size of the dots represents the mean expression of the marker across the datasets. Uppercase genes are from human datasets, while the others are from mouse ones. The outcomes of this analysis are interesting. First of all, it appears there is not a clear and coherent trend across the different datasets. Some genes exhibit a coherent trend across datasets, while others appear to be more scattered. For example, the gene *Adarb2* consistently appears in the top right corner, indicating high specificity and fraction expression (which is the most optimal case). This shows a high consistency throughout the different datasets and organisms, with an outstanding uniformity in the high values for both metrics. Hence the *Adarb2* gene is an optimal marker for the CGE-derived interneurons. At the opposite end of the spectrum, we have *Htr3a*, which is the only purely immunohistochemistry-derived marker in our list. One can immediately notice how this gene is grouped overall in the bottom right. This represents a high specificity but a low fraction expression. It means that only a few cells express it, despite being in the specific cluster. With this example, we highlight a case where it is clear that investigating only one of the two metrics could be misleading. This analysis clearly shows that *Htr3a* is not an optimal marker, despite being a CGE-derived cells marker that the immunohistochemistry studies commonly propose. In general, there are no other markers with, particularly consistent performances. Nonetheless, we can make some observations.

**Fig. 3.**
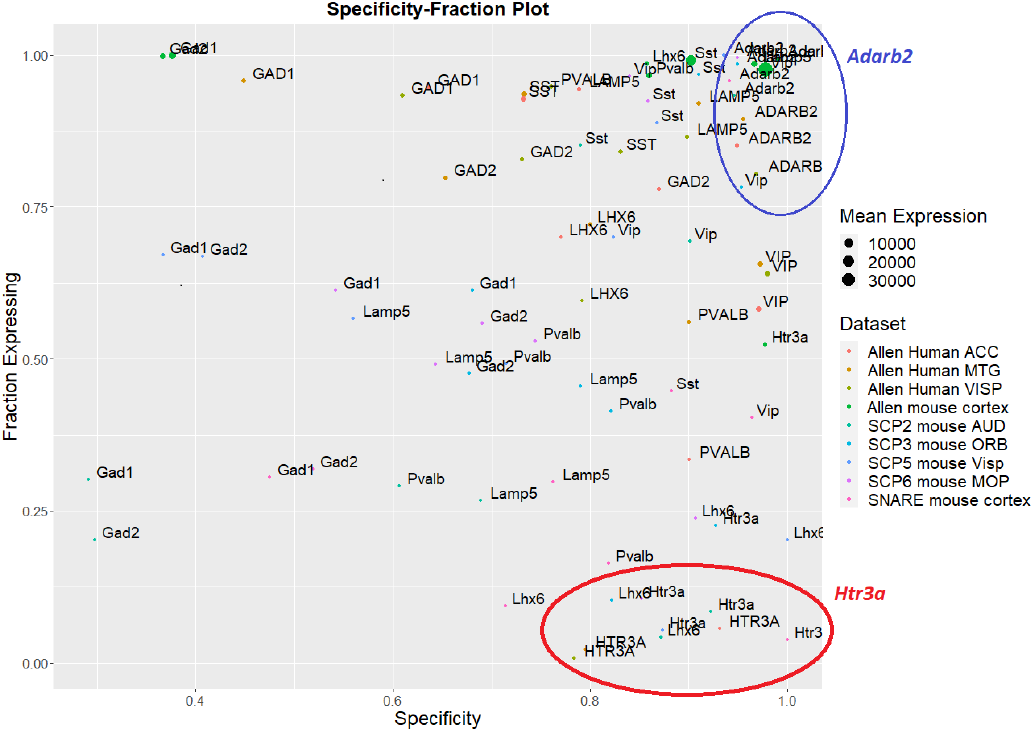
Specificity-Fraction expressing plot. Each dot is a marker from a dataset indicated by the color. The size is directly dependent to the mean expression. We can see that markers like *Adarb2* have similar high performance between datasets, indicating strong overall marker reliability. The opposite case is *Htr3a* which always has low fraction expressing, meaning that only a few cells express it.

In Figure 4, we plot the average point for each marker across the different datasets. First of all, the *Sst* and *Vip* genes have an overall high performance. It is not as high and consistent as in the *Adarb2* case. However, it shows how these genes are suitable for identifying the related cell types. Other genes (like *Pvalb, Lamp5,* and *Lhx6)* appear to have more variance in performance with an average behavior that is not particularly strong as the previous ones. Interestingly *Gad1* and *Gad2* seem to have lower performances. Specifically, they have a low *specificity,* even though in some samples, they have very high *fraction expressing* (up to 1). This is probably a consequence of how we computed the metrics for them. We performed all the calculations considering all datasets, while, for the remaining ones, we considered only the interneurons. Consequently, it is more probable to have other cells expressing these genes, lowering their *specificity*. It is particularly true for *Gad1*, which from further analysis appears to be frequently expressed by oligodendrocytes. This also explains why *Gad2* has consistently higher *specificity*.

**Fig. 4.**
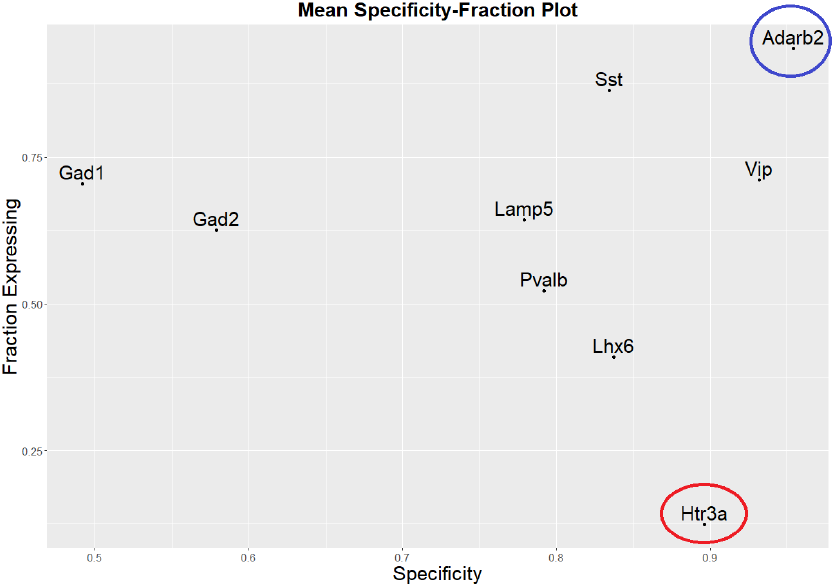
The plot of the mean performance value over datasets for each marker.

Moreover, looking at the general results, one can see how the *specificity* is consistently over 0.6, meaning the chosen set of markers is representative of the related cell types. However, the *fraction expressing* does not often reach 0.5. Therefore, the main problem is that these genes are not consistently expressed in clusters or, at least, are not detected. It is particularly problematic when one wants to label each cell by expression inspection. This is essential for rare cell types identification studies since it is crucial to label each cell correctly and not rely only on unsupervised clustering results.

From a first analysis, it appears that IR and IG have similar trends and, therefore we can use them interchangeably. For this reason, we employ only the IG and plot it with the markers score (as a medium of *specificity* and *fraction expressing*) to better assess if the two sets of metrics agree (Figure 5). The plot shows a more or less linear trend, meaning that the two sets of metrics all agree on marker performance.

**Fig. 5.**
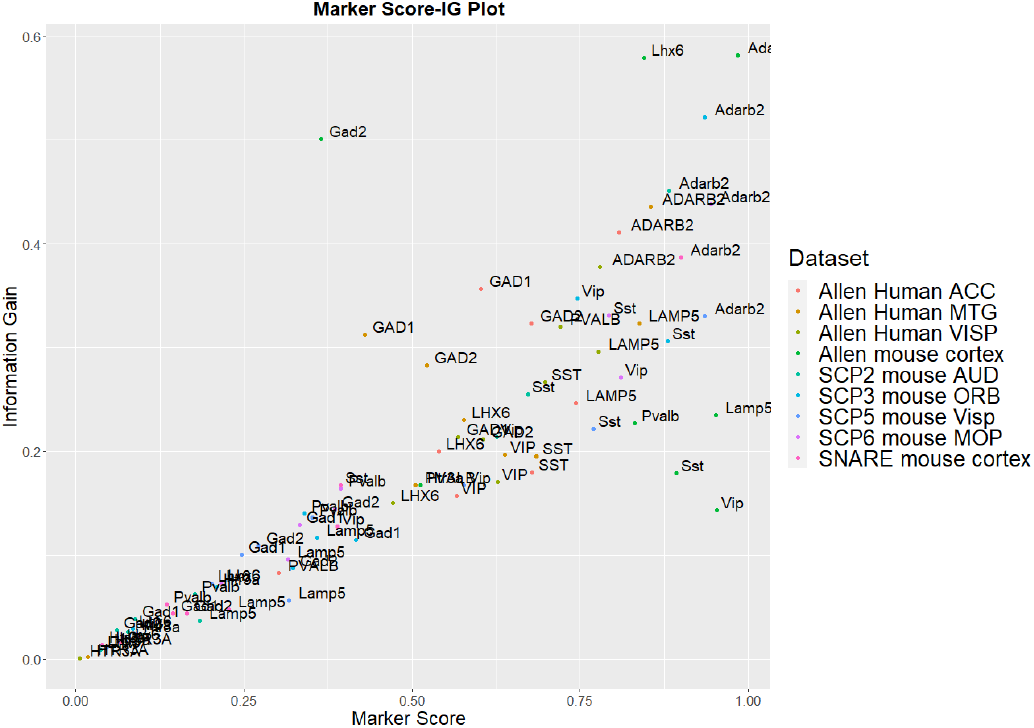
Marker Score-Information Gain plot. Each dot is a marker from a dataset indicated by the color. It shows the overall concordance of the two sets of metrics.

Concerning the possible differences within datasets, we can highlight two things. First of all, it does not appear to exist a particular relation between marker performances and the organism. This is expected since our considerations on cortical cellular heterogeneity were based on the general mammalian cortex, and, in general, human, and mouse neuronal populations are comparable [26][10]. The most prominent observation is linked to the gene’s mean expression. If we examine Figure 3, where the size of the dots is proportional to the mean gene expression, we can notice that datasets with higher mean expression values tend to have higher marker performance. This is remarkably true for the Allen mouse cortex dataset. This dataset has overwhelming higher mean expression values than the other datasets, and also the metrics for all the genes have higher values. What does it mean? Higher quality scRNA-seq experiments tend to have better catch rates, meaning they can detect and sequence more RNA molecules, resulting in a higher resolution of the gene expression inside each cell. We can appreciate it employing the marker score again. We plot the markers on the Marker Score-Mean Expression graph (Figure 6). Since the expression values vary extensively within datasets, we plot the logarithmic value of them. The resulted graph does not show a clear relationship, but a general trend is evident. The points from datasets with higher expression values tend to have also higher marker scores. However, the markers we highlighted to have better and worst performances do not seem to be influenced by this parameter. In fact, it is clear how, for example, the *Adarb2* (or *Htr3a*) marker score is high (respectively low) regardless of expression values. This indicates that the markers have general performances but have stronger reliability when employed on more specific experimental datasets.

**Fig. 6.**
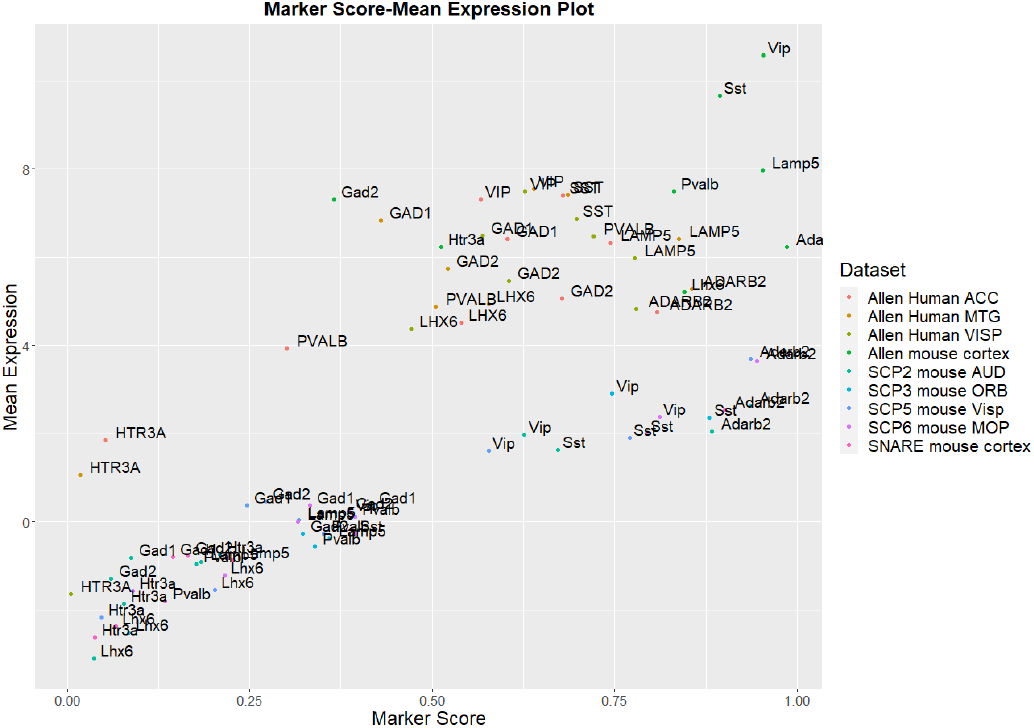
Marker Score-Mean Expression plot showing the relation between overall marker performance and the log value of the mean expression. One can see two blocks. One where the markers with low mean expression also have low marker scores, and vice versa a second one where markers with higher expression values also have higher mean scores. This show that dataset with higher mean expression result in markers with better performances.

## IV. Discussion and Conclusion

The results presented in the previous sections are non-trivial to analyze. First of all, we see that the general performance of the different markers are not all equal. For example, we noticed the marker *Adarb2*, which has optimal performance on all the metrics independently of the dataset. This shows the ability of the marker to discriminate the related cell type from the other cells, highlighting in this way his marker strength. Therefore this suggests his employment for similar works. Generally, one can expect that a marker from taxonomy studies performed on scRNA-seq studies will perform better on our datasets because they derive from the same type of experimental analysis. The opposite case is *Htr3a,* an immunohistochemistry-derived marker for the same cell type as *Adarb2.* We observe a worrying trend with *Htr3a* that cannot be considered an optimal marker, despite the literature consistently proposes it as a CGE-derived cell marker. Immunohistochemistry and transcriptional analysis are different points of view on the cell mechanisms, but there should be some level of agreement. This highlights a common problem of scRNA-seq analysis where the markers employed to label the clusters come from the immunohistochemistry literature, even if they are not always optimal for transcriptomic data. Apart from the above-mentioned markers, which have high consistency within datasets, the other genes appear to vary a lot. Our analysis remarked a correlation between the marker performance and its mean expression throughout the dataset. This is particularly interesting since it indicates that datasets with higher experimental mean expression influence the classification reliability of the markers. Therefore, we can say that if one is interested in rare cell type investigation (or, more in general, precise cell type labeling), he should employ datasets with a higher average mean expression, which ensures higher markers performances.

In conclusion, this work shows how the formal analysis of markers’ consistency and reliability supports a well-informed choice of markers for cellular heterogeneity studies. In our view this should become a habitual practice in the common single cell analysis workflow.

